# Experience improves navigational knowledge of dumpers in desert ants (*Melophorus bagoti*)

**DOI:** 10.1101/2024.07.14.603463

**Authors:** Ken Cheng, Sudhakar Deeti

**Affiliations:** School of Natural Sciences, Macquarie University, Sydney, NSW 2109, Australia

**Keywords:** Insect Behaviour, Ant Colony, Waste Management, Naive ants, Experienced dumpers

## Abstract

The Australian red honey ant, *Melophorus bagoti*, is an excellent desert navigator, performing all the activities outside the nest during the hottest periods of summer days. This species relies heavily on path integration and landmark cues for outbound and inbound navigation. Although the species navigational behaviours have been much studied, the spatial knowledge of workers that dump waste has not been investigated. In our study, we investigated the navigational knowledge of both naive and experienced dumpers by displacing them 2 metres away from the nest. Naive dumpers were not oriented towards the nest in their initial trajectory at any of the test locations, whereas experienced dumpers were significantly oriented towards the nest at all test locations. Naive dumpers were nest-oriented as a group, however, at the test location nearest to where they dumped their waste. Compared with experienced dumpers, the paths of naive dumpers were more sinuous, and naive dumpers scanned more on tests. Overall, our findings suggest that dumpers gain greater spatial knowledge through repeated dumping runs outside the nest, contributing to successful homing behaviour.

## Introduction

The eusocial ants feature specialisation of labour in their societies (Hölldobler and Wilson, 1990). Specialisation is found even when workers are monomorphic and look similar. Different ants assume different roles, flexibly to an extent based on colony demands. In the desert ants *Cataglyphis* genus, individuals may perform different roles such as foraging and brood care responsibilities depending on environmental conditions (Wehner et al., 1983). In leafcutter ant colonies, similar-looking workers specialize in cutting and transporting leaf fragments or tending the fungus gardens (Hölldobler and Wilson, 1990; Weber, 1972). In army ant colonies, workers have a uniform appearance but perform various tasks, with certain workers foraging and others maintaining and protecting the temporary bivouac nests during colony migration (Franks, 1989). Specialisation among similar-looking workers also applies to our current study species, the Australian red honey ant, *Melophorus bagoti* Lubbock, 1883. Different workers carry out three outdoor tasks that we have identified in research on this species: foraging, excavating the nest, and dumping waste. Foragers scavenge for dead arthropods in the heat of the day during summer months and also bring home plant materials, including sap (Muser et al., 2005; Schultheiss and Nooten, 2013). The navigational strategies with which they accomplish this task has been much studied (Cheng et al., 2009; Cheng et al., 2014; Wehner, 2020). Excavators maintain the nest, mostly by carrying excess sand out of the nest and dropping it nearby (Deeti et al., 2023a; Deeti et al., submitted a). This account concerns dumpers, which carry waste material such as leftover food and dead nestmates out of the nest, taking such stuff some distance before tossing it with a characteristic lunge (Deeti et al., 2023; Deeti et al., submitted b).

Waste management, in which dumping is a key job, serves an important function in keeping pathogens and potential pathogens away from a crowded nest, in which the spread of disease can be costly or even deadly (Cremer et al., 2007; Cremer et al., 2018). The workers collect scavenged food and nest materials while discarding items deemed unnecessary (Hart and Ratnieks, 2002; Czaczkes et al., 2015). Given that waste may harbour pathogens, rapid disposal becomes paramount for maintaining colony health and hygiene (Cremer, 2019). The red honey ant *M. bagoti* employs two distinct strategies. During the summer months, they transport waste materials away from the nest, while during the winter hibernation, they deposit waste in subterranean latrine channels/tunnels, approximately 10 to 12 cm below ground level, a behaviour likely persistent throughout the year (Deeti et al., 2023a). Dumpers show a kind of ‘pathogen sensitivity’ in dumping (Deeti et al., 2023a). The effort expended increases with the potential pathogenicity of the waste. Waste posing a higher risk of harbouring pathogens is transported farther away from the nest compared to harmless materials. Potentially pathogenic items dropped experimentally into the nest, including dead ants, food waste, bits of dead moth, and cicada shells, were dumped at average distances exceeding 5 metres, while items with low pathogenic potential were dropped at average distances less than 1.2 metres.

For all these outdoor workers to continue in their roles, they must return to the nest at the end of each task. Foragers of this species take multiple *learning walks* around their nest, 3–7 in number, before heading off on a foraging trip (Deeti and Cheng, 2021a), as do other species of ants (*Cataglyphis fortis* and *noda*: Fleischmann et al., 2016; Fleischmann et al., 2018; *Myrmecia croslandi*: Jayatilaka et al., 2018; reviews: Freas et al., 2019; Zeil and Fleischmann, 2019). Ants walk in loops around their nest, canvassing all directions and stopping occasionally to pause and look in different directions, including, importantly, the nest direction. Over multiple walks, the loops get bigger, and the learner ventures farther from the nest. Recent work revealed that *M. bagoti* excavators also take a learning walk, a single one, before taking on excavating duties, even though the unwanted sand is only jettisoned a maximum of ∼15 cm from the nest entrance (Deeti et al., submitted a). The single learning walk resembles the first learning walk of foragers. Such walks are presumed to help outdoor workers learn their visual surround to aid navigation on their trips. Indeed, in *M. bagoti*, a single learning walk enables them to head in the home direction from an arbitrary location 2 m from the nest, although not from 4 m (Deeti and Cheng, 2021a; Deeti et al., submitted a). In one nest whose ‘pre-job’ behaviour has been documented, however, dumpers did not take any learning walks before their first dumping job (Deeti et al., submitted b). This finding surprised the authors because the journey encompassed 3 or even, in one case, 4 metres of travel distance from the nest. Nevertheless, although their navigation was less efficient than those of experienced dumpers, the first-time dumpers managed to get home, seeming to have learned on the job.

The limited extant data on dumping behaviour in ants leave key open questions to be examined. A chief open question is the visual knowledge of the naive dumpers. Can they home from a range of directions near the nest? To test such knowledge of the visual surrounds, the standard and accepted method is to give displacement tests to returning ants that have nearly reached to their nest. On an outbound job, whether foraging, excavating, or dumping, desert ants can call on multiple strategies in their navigational toolkit. They can use path integration (PI): to continuously track the distance and compass direction from their starting point (the nest) of all path segments during the trip (Mittelstaedt and Mittelstaedt, 1980; Mittelstaedt and Mittelstaedt, 1982; Müller and Wehner, 1988; in *M. bagoti*: Narendra 2007; Cheng et al., 2009). PI enables an animal to orient and face in the direction of the starting point without any knowledge of the nest’s visual surroundings. If an ant has just about returned to its nest, however, its PI vector is near 0 and is not informative as to a direction of travel. Displacement tests force an ant to use another strategy in the toolkit, view-based navigation (Cheng et al., 2009; Graham and Cheng, 2009a; Graham and Cheng, 2009b; Wystach et al., 2011; Deeti et al., 2020; Wehner 2020). Views around the nest are learned and later used to navigate home.

In this study, we examined the navigational capabilities of both naive (inexperienced) and experienced *M. bagoti* dumpers using displacement tests. Both experienced and naive dumpers were tested at locations 2 m from the nest in four cardinal directions, with one of those directions being familiar to naive ants because their dumping trip was in that sector. We videotaped the start of these tests and documented the characteristics of dumpers’ behaviours, including scanning behaviour, travel speed, and gaze-oscillation velocity, comparing naive and experienced dumpers. As before with characteristics of travel on the dumping job (Deeti et al., submitted b), we predict significant differences between the two groups. On tests, we also expected experienced dumpers to be well oriented in the home direction at all test sites, while naive dumpers would only be well oriented in the test location nearest their dumping spot.

## METHODS

In this study, we investigated the navigational knowledge of experienced and naive dumpers. We conducted the tests on one *Melophorus bagoti* nest (Lubbock, 1883) between November 2023 and February 2024. The nest was situated close to the Centre for Appropriate Technology, located 10 km south of Alice Springs, NT, Australia (23°45′28.12″S, 133°52′59.77″E). The surrounding habitat of the nest was an open space of a semi-arid desert, scattered with a mix of *Acacia* bushes, buffel grass (*Pennisetum cenchroides*) and *Eucalyptus* trees (Deeti and Cheng 2021b; Deeti et al., 2020), forming a distinct visual panorama. In Australia, there are no specific ethical guidelines pertaining to ant studies, and our experimental procedures were entirely non-invasive.

### Experimental animals

The red honey ant, *M. bagoti*, is the most thermophilic ant species found on the Australian continent (Christian and Morton, 1992). We observed that ants were usually active outside the nest when the temperature ranged between 32 and 40°C. During the hot southern summer, these *M. bagoti* ants engage in foraging. We have noticed that the foragers scavenge mainly dead spiders, sugar ants, termites, centipedes, moths and other dead arthropods. They also collect sugary plant exudates and seeds (Muser et al., 2005; Schultheiss and Nooten, 2013). The *M. bagoti* foragers typically forage solitarily, covering distances up to 50 m from the nest (personal observations), relying on path integration and terrestrial visual landmarks rather than chemical trails to find their way to the nest (Cheng et al., 2009). Some workers, called dumpers, are responsible for transporting and dumping waste material, such as unwanted food and dead ants, from the nest to more than 8 meters away from the nest (Deeti et al., 2023a).

We tested naïve and experienced *M. bagoti* dumpers. Considering the average foraging span of *M. bagoti* ants outside the nest is estimated at five days (Muser et al., 2005), we painted all the emerging ants using a black dot on their abdomen for nine consecutive days. Beyond this period, any unpainted ants emerging with waste from the nest were identified as naive dumpers, whereas black-painted ants were identified as experienced dumpers. For the naive group, we tested only dumpers that did not engage in a learning walk before going on a dumping trip, so that they were completely naive to the visual surroundings. We tested N=15 ants for both experienced and naive dumpers. We found four naive ants performing learning walks that later engaged in dumping. We painted these four a different colour to exclude them from the study. From that lot, we noted that three naive dumpers performed two learning walks and one performed a single learning walk before starting the dumping activity.

### Experimental procedure

On the initial day of the study, we cleared the vegetation surrounding the tested *M. bagoti* nest within a radius of 10 m to improve the visibility of the dumpers’ activity. To enhance the visibility of the red ants against the red soil background, we spread fine white sand within the recording area at the displacement sites. The four test locations were situated 2 m from the nest towards North, East, South and West (2mN, 2mE, 2mS, 2mW).

We followed dumpers during their outbound trip carrying nest-related wastes and inbound trips returning to the nest after they dumped nest-related wastes. We captured the inbound dumpers within 10 cm of the nest and displaced them to four test locations. We tested each dumper at these four locations in a random order. Once the tests were completed, we marked the tested dumper with a yellow colour on the abdomen to avoid repeated testing, and then we released the tested ant back into the nest.

### Tracking and Data extraction

In this study, we analysed the ants’ orientation and path characteristics at the displacement sites. To record dumpers’ path, we used a Sony Handy camera (FDR-AX700, 25 fps, 3860 x 2160 pixels) positioned on a tripod at a height of 1.2 metres from the ground, recording an area of 1 × 1 metre centred at the displacement location. We used the animal tracking program DLTdv8 (version 8.2.9) in MATLAB (2022B) to extract frame-by-frame coordinates of the dumper’s head and thorax — specifically the tip of the head and the middle of the thorax (see Fig. S1). During displacement tests, ants frequently displayed a series of stereotypical successive fixations in different directions by rotating on the spot at one location, known as a “scanning bout” (Deeti et al. 2023b), and we extracted the number and duration of scanning bouts.

### Data analysis

We used the extracted frame-by-frame coordinates to calculate the final heading direction, speed, gaze direction, gaze angular velocity and path characteristics. We determined the final heading direction by the thorax–head direction as an ant exited the recording area. This vector represented the orientation of each ant at that specific moment. To understand how quickly and directly the ants moved at each displacement location, we calculated the speed of the workers along with several characteristics of the trajectory. These path characteristics were based on the positions of the thorax across frames. Speed refers to the magnitude of an ant’s velocity and was calculated as the average over the entire trajectory for each ant, excluding the stopping durations. We measured the speed at which ants change their gaze direction by observing how quickly their gaze angle, defined by the thorax–head vector, changes over time. As ants walk, their gaze constantly shifts.

To understand the path characteristics at the different displacement-test locations, we used three indices of straightness: *path straightness, sinuosity*, and 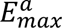, each of which relates to the directness of navigation towards a destination. *Straightness* is computed as the ratio of the straight-line distance between the release point at the displacement site and the final point in the frame before the ant moved out of the recording area to the overall length of the path (Batschelet, 1981; Islam et al., 2021; Islam et al., 2023; Deeti et al., 2023c). *Straightness* ranges from 0 to 1, with larger values indicating straighter paths, while smaller values indicate more curved or convoluted paths. *Sinuosity* is an estimate of the tortuosity in a path, calculated as 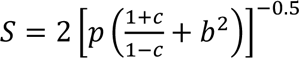 where *p* is the mean step length, *c* is the mean cosine of turning angles and *b* is the coefficient of variation of the step length. A trajectory *step* is the movement between the positions of the animal (thorax positions) recorded in consecutive video frames. Accordingly, step lengths are the Euclidean distances between consecutive points along a path, and turning angle refers to the change in direction between two consecutive steps. Sinuosity varies between 0 (straight) and 1 (extremely curved) (Benhamou, 2004; Lionetti et al., 2023). The maximum expected displacement of a path, 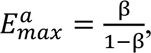, where β is the mean cosine of turning angles, is a dimensionless value expressed as a function of the number of steps and is consistent with the intuitive meaning of straightness (Cheung et al. 2007). Larger maximum expected displacement values indicate straighter paths, hence greater displacement, while a smaller value suggests more localised or constrained movement. Paths were characterized and visualized in R (version 4.2.1; R Core Team, 2020) using the packages trajr (McLean and Skowron Volponi 2018) and Durga (Khan and McLean 2023).

We compared the panoramic images from each test location to the nest by computing the rotational image difference function (rotIDF) between the nest panorama and each test-location panorama (Zeil, 2012). The pixel differences were calculated for each 1° shift in the test-location panorama, following methodologies outlined in previous studies (Islam et al., 2022). We used Richo Theta cameras to obtain panoramic images from each test location and the nest.

### Statistical analysis

We performed a Rayleigh’s test to assess whether the distribution of headings for each condition was uniform (*p* > 0.05). For the naive ants, we tested an additional heading distribution consisting of the test performance of each ant at the test site nearest to where the ant dumped its waste. This distribution was not compared with any other distribution. For the other distributions, in case two distributions turned out non-uniform, we planned to compare the mean direction of the two groups using the Watson–Williams test (alpha = 0.05). Additionally, we examined if final heading orientations significantly clustered around the nest direction at 0 degrees by checking whether 0 degrees fell within the 95% confidence interval (CI) of orientations (Watson tests). V-tests were also conducted, with alpha set at alpha = 0.05, to determine if the mean headings were notably clustered around the nest direction. Single-sample log likelihood ratio tests were also conducted to investigate whether the heading distributions of the ants were uniform in each test condition.

We used generalised linear mixed models to investigate whether the displacement location affected dumpers’ *path straightness*, *sinuosity*, 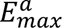, speed and gaze angular velocity. The model treated dumper type (experienced or naïve), as a between-subjects factor and displacement locations as a within-subject variable that predict the dependent variables. The LM was formulated using the lme4 package (version 1.1-27) and fitted using the lmer function (Bates et al., 2015). Since *path straightness* and *sinuosity* are bounded measures (0– 1), we used the binomial family, whereas all of our other response variables are only bounded by zero, and so for them, we applied the Gaussian family of models. We used the Tukey post hoc test to perform pair-wise post hoc comparisons for each of our models. Since we tested multiple dependent variables computed from the same data sets of trajectories on tests, we adopted alpha = 0.01 to lower type 1 errors. Statistical analyses were conducted using R (version 4.3.1).

## RESULTS

The majority of naive dumpers did not engage a learning walk before heading off on their first dumping job: 15 did not while 4 did. Of the latter, 3 took 2 learning walks while one took a single learning walk. As indicated in the Methods, we excluded the entire latter cohort and studied only the naive dumpers that did not do a learning walk. Our informal observations on experienced dumpers suggested that most of these workers stuck to their job, undergoing a period of intensive dumping activities typically spanning 3 to 4 days. They would have had taken many trips of > 5 m in distance (based on earlier data in Deeti et al., 2023a) when they were tested in this study. None were observed to take learning walks.

In order to understand the navigational knowledge of these experienced and naive dumpers, after their dumping activity, each ant was captured just before it entered the nest and displaced to 2 m from the nest in four different locations to the North, East, South and West. The ants’ final headings at each location showed that the experienced dumpers in the 2-m displacements were oriented towards the nest direction of 0 deg from all four cardinal directions. In contrast, naive dumpers that were displaced 2 m from the nest were not oriented towards the nest direction at any release point (Fig. 1). By the Rayleigh test, the experienced dumpers’ initial orientations were non-uniformly distributed in all 2-m displacement tests whereas the naive dumpers were scattered, with uniformly distributed headings (Table 1, Fig. 1). In addition, experienced dumpers in 2m displacement conditions showed significant V-test results in the nest direction, and the means of their 95% confidence interval of initial heading values include the nest direction 0 deg (Watson test, *p* > 0.05). The log likelihood ratio test for the experienced dumpers on 2m tests failed to reject the hypothesis that the distribution was clustered in the home direction (*p* ≥ 0 .05, k ≥ 0, χ2 ≤1). For the naive dumpers on 2m displacement tests, however, the log likelihood ratio rejected the hypothesis that the mean value of distribution was equal to the predicted value (home direction) (*p* = 0.0006, k = 0.64 and χ2=5.6), meaning that the headings were oriented in a different direction from the nest direction. We checked whether the naive dumpers were well oriented at the test location nearest their dumping point. The majority of the naive dumpers oriented in the general direction of the nest (Table 1 Fig,1I). Both the Rayleigh test and the V-test returned significant values, indicating an oriented distribution in the nest direction.

**Fig. 1.**
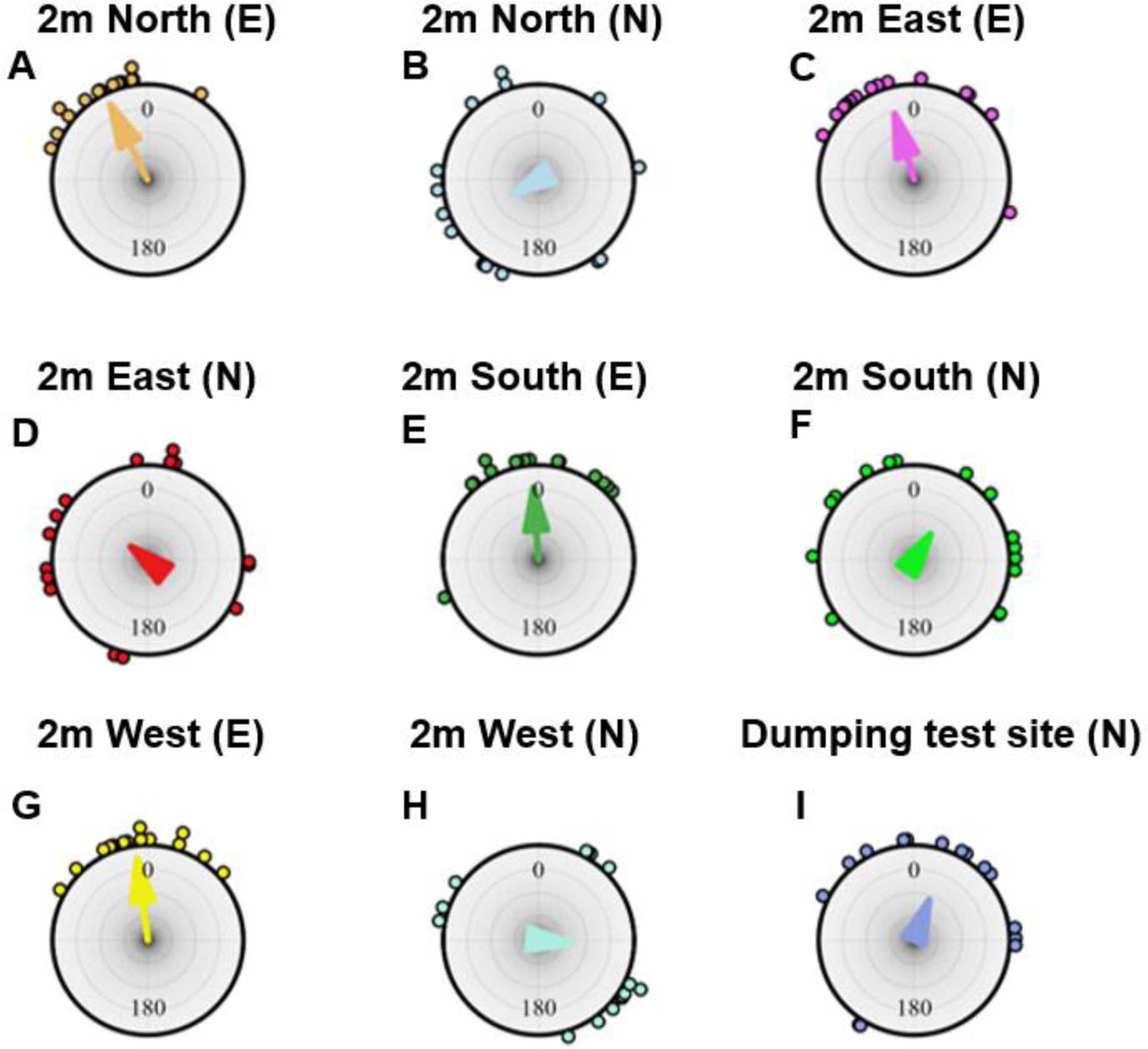
Headings on displacement tests. Circular histograms of initial headings of foragers during the displacement tests at 2m North in experienced (A) and naive (B) ants, at 2m East in experienced (C) and naive (D) ants, at 2m South in experienced (E) and naive (F) ants, and at 2m West in experienced (G) and naive (H), and at the test site nearest the dumping location of each naive ant (I). In the figure title E denotes experienced and N denotes naive. In the histograms, the nest direction is set at 0°. The arrows denote the length and direction of the mean vector.

**Table 1:** Statistical results for initial heading directions of experienced (E) and naive (N) dumpers at 2m North, 2m East, 2m South and 2m West displacement tests. The last line concerns naive dumpers at the test site nearest to where they dumped their waste.

| Test | <u>Mean vector</u> | <u>95% confidence interval</u> |  | <u>Rayleigh test</u> |  | <u>V test detection 0<sup>0</sup></u> |  |
| --- | --- | --- | --- | --- | --- | --- | --- |
| | $\mu$ | Minus | Plus | Z | P | Z | P |
| 2m North (E) | 333.71° | 321.26° | 346.16° | 12.91 | < 0.0007 | 4.557 | < 0.0002 |
| 2m North (N) | 227.94° | 179.31° | 276.53° | 2.81 | 0.057 | -1.59 | 0.944 |
| 2m East (E) | 345.10° | 321.82° | 8.388° | 8.8 | < 0.0002 | 4.055 | < 0.0005 |
| 2m East (N) | 328.47° | 164.2° | 295.88° | 0.51 | 0.61 | 0.864 | 0.196 |
| 2m South (E) | 358.27° | 337.28° | 19.27° | 9.72 | < 0.0005 | 4.409 | < 0.0004 |
| 2m South (N) | 25.69° | 312.211° | 99.17° | 1.55 | 0.214 | 1.589 | 0.056 |
| 2m West (E) | 359.92° | 341.572° | 18.29° | 10.82 | < 0.0001 | 4.652 | < 0.0001 |
| 2m West (N) | 112.04° | 31.736° | 192.35° | 1.38 | 0.254 | -0.625 | 0.732 |
| Dumpers at nearest<br>test site (N) | 28.69° | 314.211° | 89.17° | 4.82 | 0.0214 | 2.89 | <0.005 |

We checked whether experienced dumpers differed from naive dumpers in their path characteristics at the four different displacement locations. Firstly, the experienced dumpers exhibited lower tortuosity in their paths, indicating that they changed their travelling direction fewer times compared to the naive dumpers, who tended not to take direct paths. The linear mixed model ANOVA showed significant differences in *sinuosity* between the experienced and naive groups (F_1, 59_ =33.64, *P*<0.0005) (Fig.2A). However, no significant difference was found within the experienced and naive groups across test locations (F_3, 57_ =0.17, *P*=0.91). Additionally, the model did not detect any significant interaction (F_3, 57_ =0.44, *P* =0.71). Overall, the sinuosity measure confirmed that the experienced dumpers showed more efficiency in paths than did the naive dumpers. Secondly, in 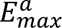, the degree of displacement varied between the experienced and naive dumpers. The linear mixed-model ANOVA showed significant differences in 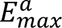 between the experienced and naive groups (F_1, 59_ =32.29, *P*<0.0005) (Fig.2A). However, no significant difference was found within the experienced and naive groups across locations (F_3, 57_ =1.2, *P*=0.29). Additionally, the model did not detect any significant interaction (F_3, 57_ =0.18, *P*=0.9). Finally, with *straightness*, the analysis of variance found statistical significance only between groups (F_1, 59_ =55.9, *P*<0.0005), with a trend across test locations (F_3, 57_ =3.52, *P*<0.018) and in the interaction (F_3, 57_ =2.79, *P*<0.04).

**Fig. 2.**
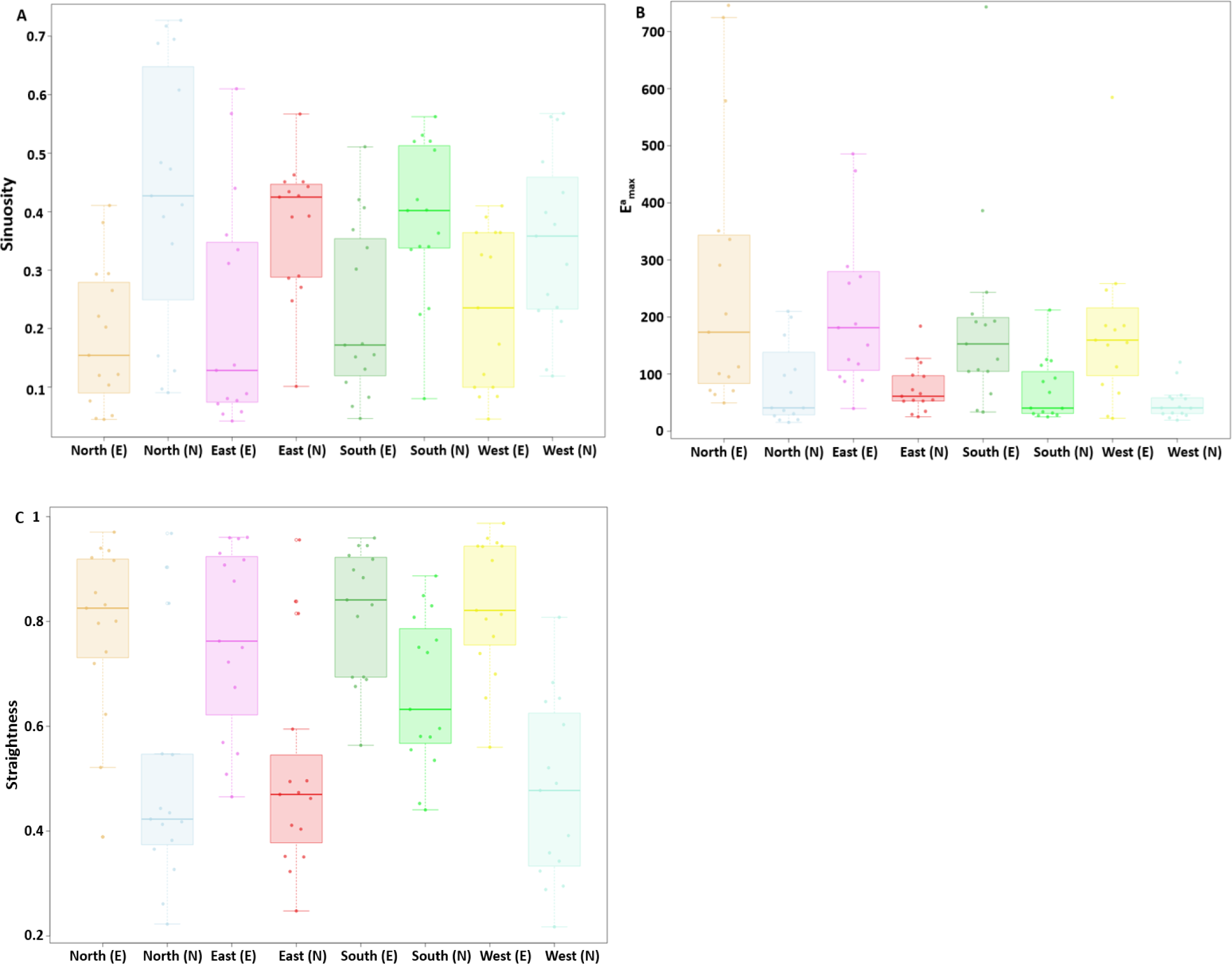
Path characteristics of dumpers at various test locations. Path characteristics of ants at different displacement locations A) *sinuosity*, B) 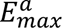 and C) *straightness*. Box plots display the median (line inside the box), interquartile range (box), and extreme values excluding outliers (whiskers). Individual data points are shown as dots. In the x axis of the figure legend, ‘E’ after the direction name denotes experienced, whereas ‘N’ denotes naive.

During the displacement test we found that the lack of experience in naive dumpers had noticeable impact on the speed of the ants (Fig. 3A and 3B). The linear mixed-model ANOVA showed significant differences in mean speed between the experienced and naive groups (F_1, 59_ =31.94, *P*<0.0005) (Fig.3A). However, no significant difference was found within the experienced and naive groups across locations (F_1, 59_ =0.82, *P*=0.48). Additionally, the model did not detect any significant interaction (F_3, 57_ =1.8, *P*=0.14). As for gaze angular velocity, the linear mixed-model ANOVA showed a trend between the experienced and naive groups (F_1, 59_ =5.5, *P*<0.02) (Fig.3B). However, no significant difference was found within the experienced and naive groups across locations (F_3, 57_ =0.82, *P*=0.47). Additionally, the model did not detect any significant interaction (F_3, 57_ =2.1, *P*=0.1).

**Fig. 3.**
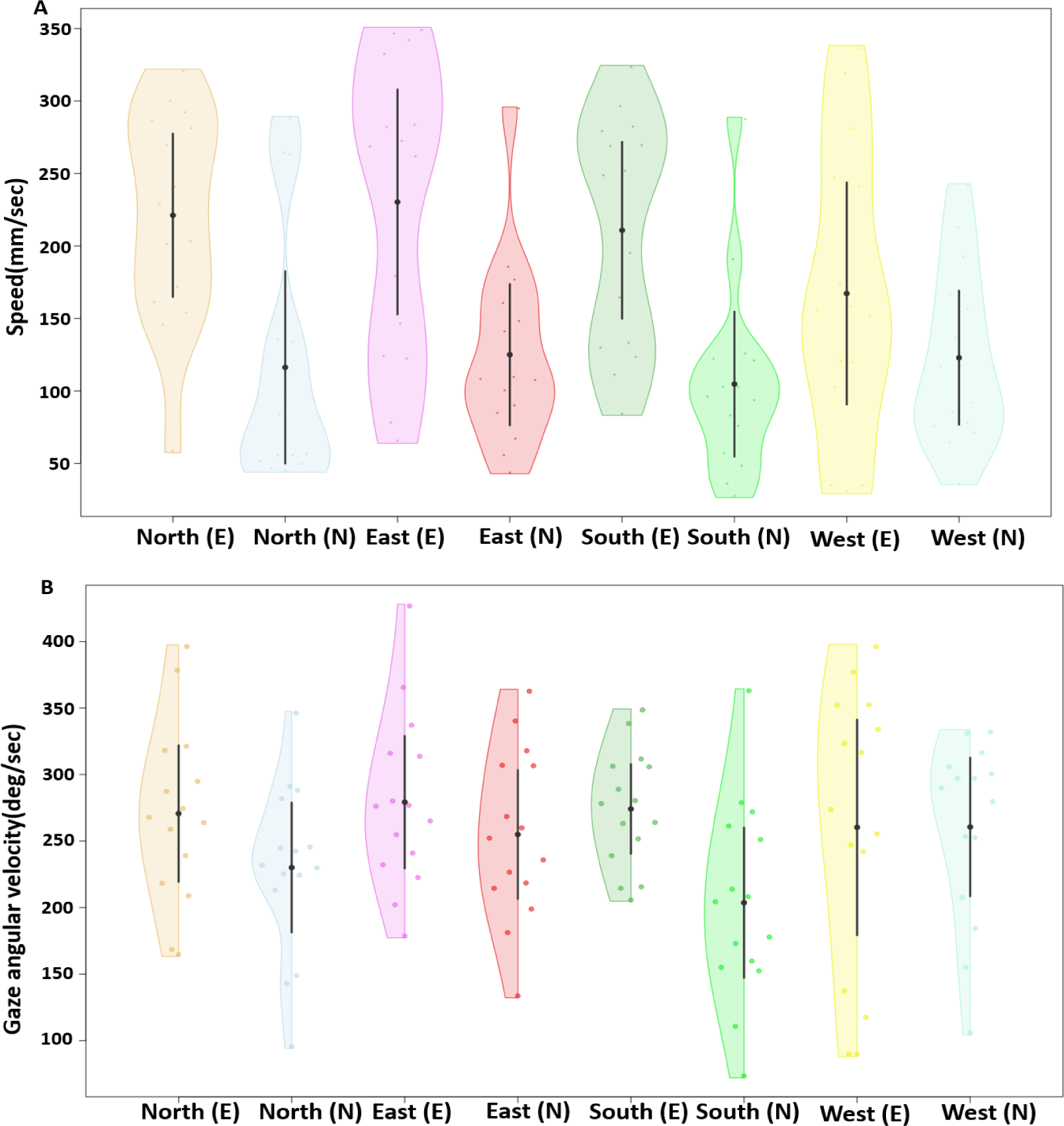
Speed and gaze angular velocity of displaced ants. Comparison of speed and gaze angular velocity of ants at different displacement locations. The violin plots show the mean speed of ants across their entire trajectory at each displacement location (A). The half-violin plots show the distribution of bootstrapped differences of mean gaze angular velocity of ants at the various displacement locations (B). In (A) and (B), the solid dot shows mean, while the vertical bar shows 95% confidence interval of the mean. In the x axis figure legend, ‘E’ after the direction name denotes experienced, whereas ‘N’ denotes naive dumpers.

We utilized rotIDFs to assess the visual similarity of each displacement location to the nest panorama (Fig. 4). Comparisons with the nest panorama revealed detectable minima for all displacement locations. The depth of these minima showed only small differences across three displacement locations to the North, East, and West, ranging from 9.4 to 11.4, with the South location having a much lower depth of only 2.1. Nevertheless, this small depth was enough for experienced dumpers to orient homewards (Fig. 1E).

**Fig. 4.**
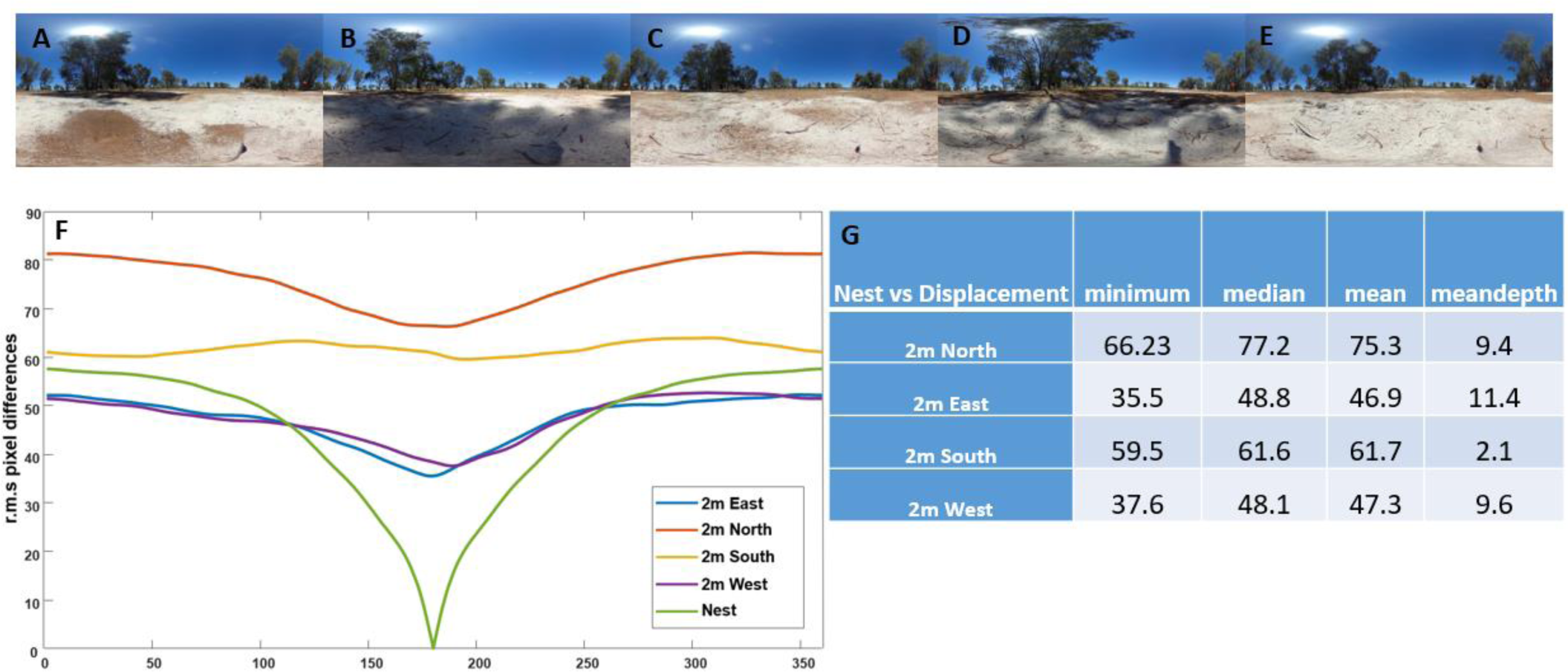
Nest Panorama and rotational image differences. A to E). Panoramic views at various locations, all aligned towards the nest direction: at A) the Nest, B) 2m North, C) 2m East, D) 2m South, and E) 2m West. F). The rotation image difference function for each location when compared with the nest panorama facing the direction of the nest. G). Table shows the minimum, median, mean and mean depth (mean-minima) of mismatch at each test location when compared with the nest panorama facing the direction of the nest.

During the displacement tests, we found that naive dumpers showed more scanning behaviour compared to experienced dumpers (Fig. 5A and 5B). The linear mixed-model ANOVA showed significant differences in mean number of scanning bouts between the experienced and naive groups (F_1, 59_ =72.95, *P*<0.0002) (Fig.5A). However, no significant difference was found within the experienced and naive groups across displacement locations (F_3, 57_ =0.74, *P*=0.52). Additionally, the model did not detect any significant interaction (F_3, 57_ =0.61, *P*=0.60). Similarly, with the scanning-bout durations, the linear mixed-model ANOVA showed significant differences between the experienced and naive groups (F_1, 59_ =9.23, *P*<0.005) (Fig.5B). Nevertheless, no significant difference was found within the experienced and naive groups across displacement locations (F_3, 57_ =0.2, *P*=0.89) and the model did not detect any significant interaction (F_3, 57_ =2.45, *P*=0.06).

**Fig. 5.**
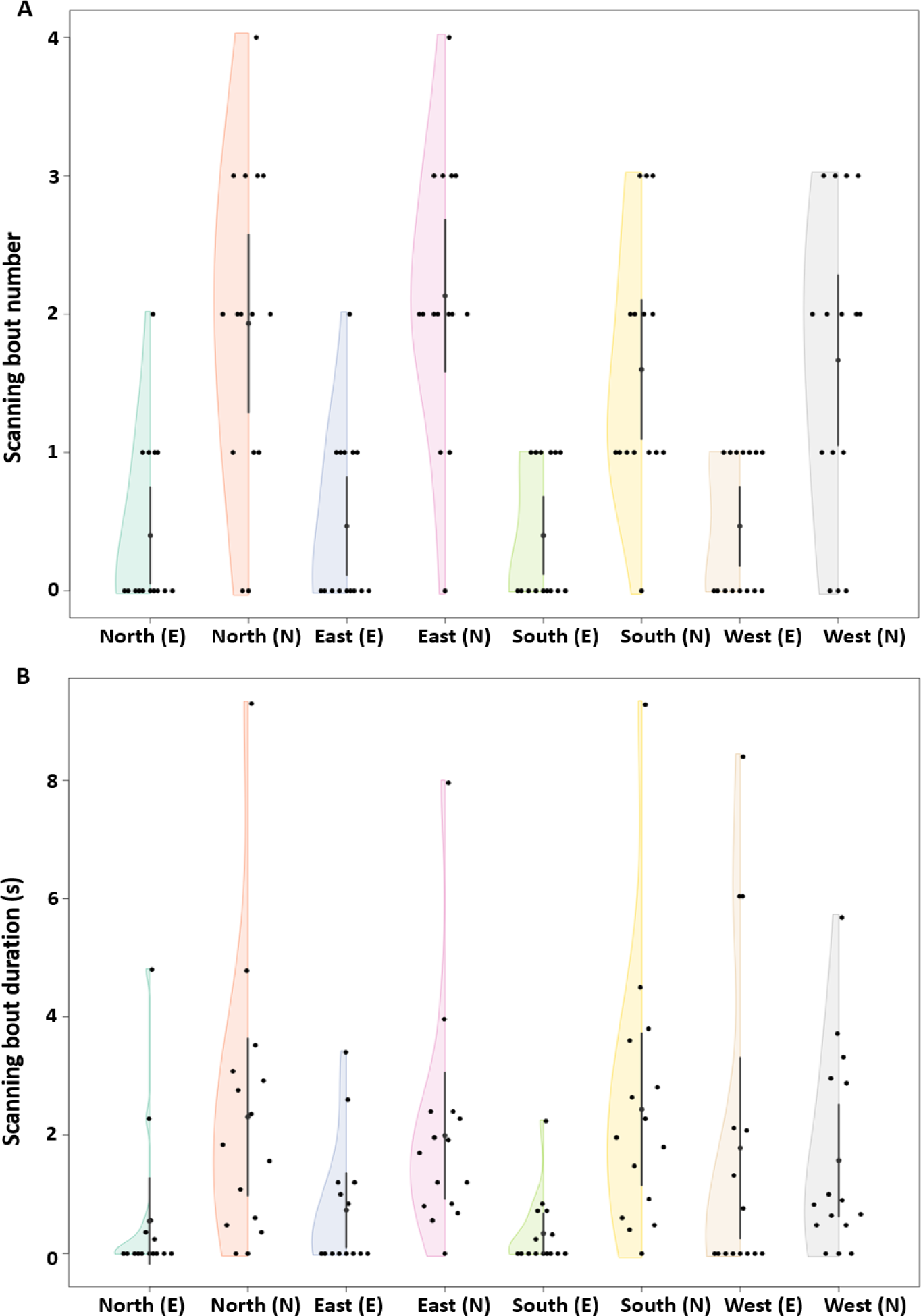
Number and timing of scanning bouts. Number of scanning bouts (A), scanning-bout duration (B), of the experienced and naive dumpers during the displacement tests. The half violins show the distribution of bootstrapped differences, the solid dot shows the mean, while the vertical bar shows 95% confidence interval of the mean. In the x axis figure legend, ‘E’ after the direction name denotes experienced, whereas ‘N’ denotes naive dumpers.

**Fig. 6.**
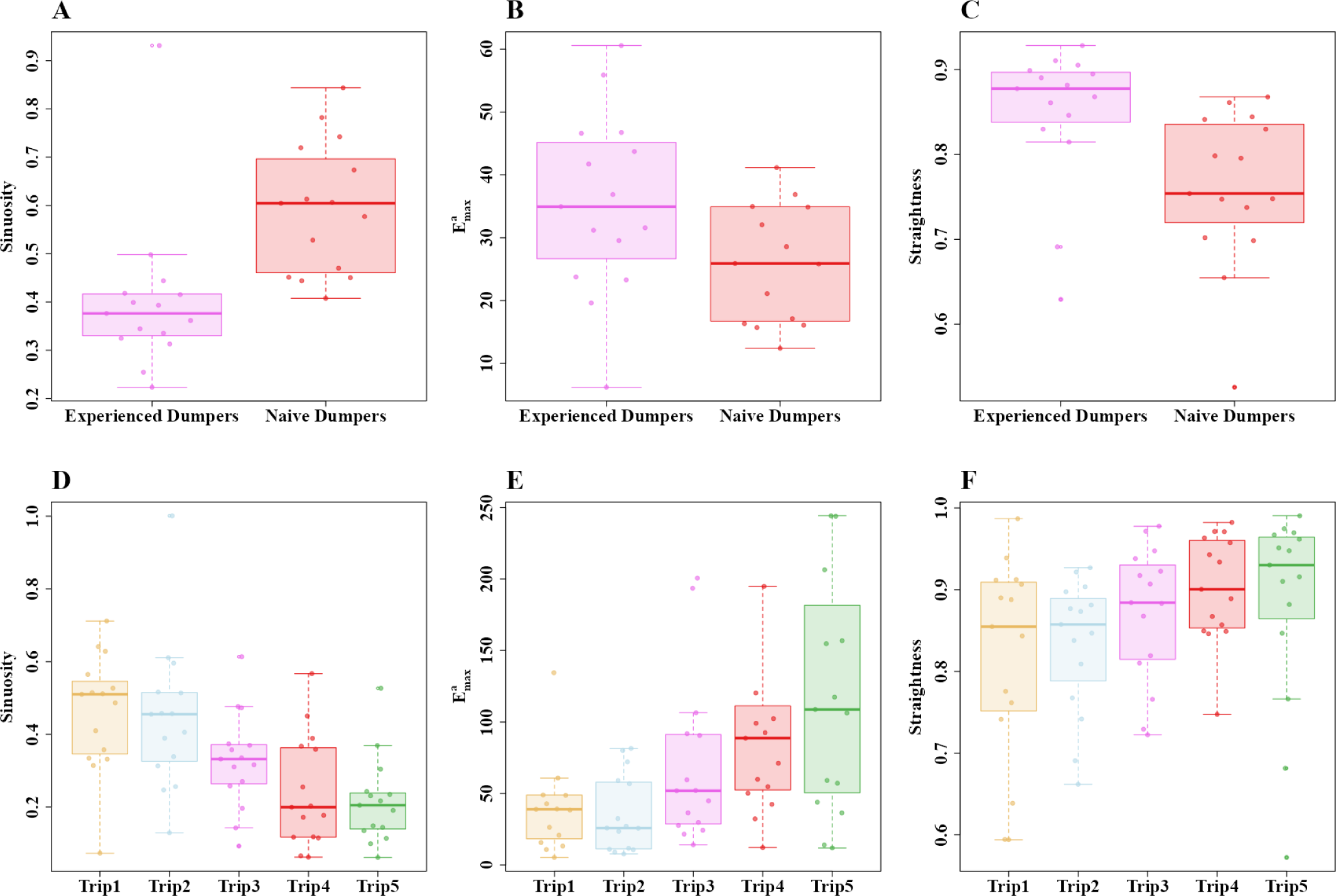
Path characteristics of experienced and naive dumpers and during the five consecutive dumping trips. A) *sinuosity*, B) *E_max_* and C) *straightness*. Box plots display the median (line inside the box), interquartile range (box), and extreme values excluding outliers. Individual data points are shown as dots.

**Fig. 7.**
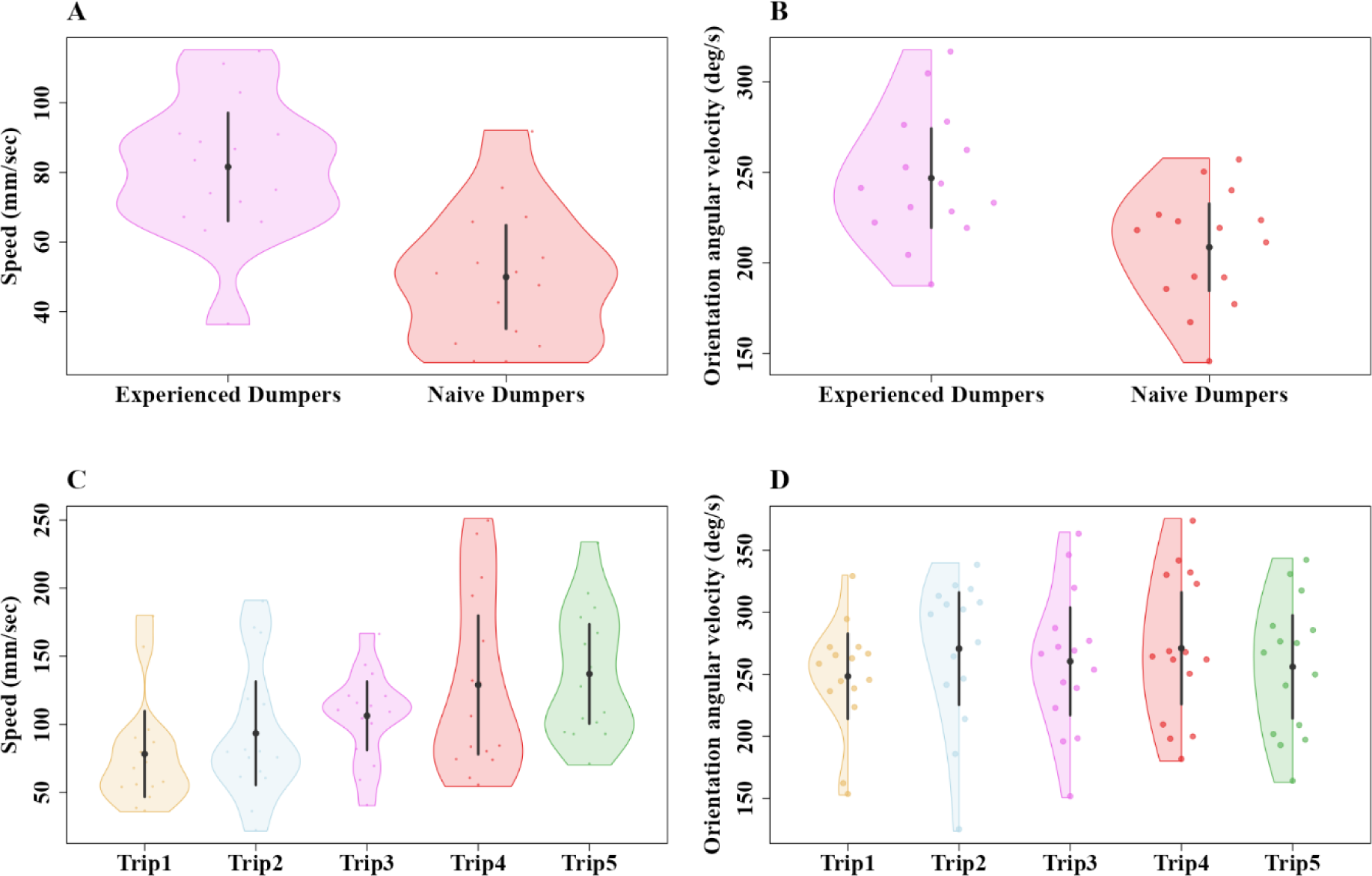
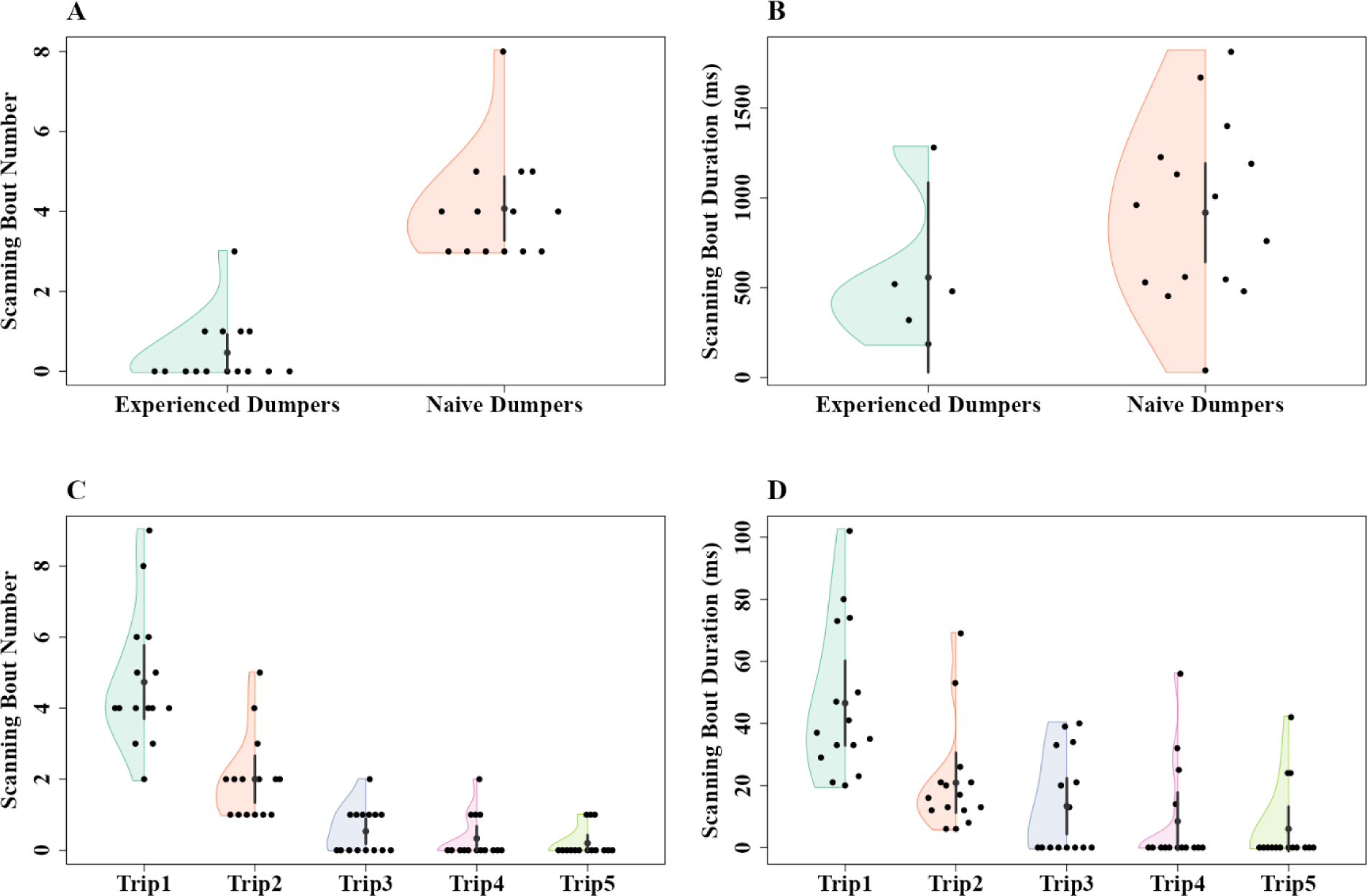
Comparison of speed and gaze angular velocity of experienced and naive dumpers and over five consecutive dumping trips. Two examples of the speed of single ants each condition plotted against time (A). The violin plot shows the mean speed of experienced and naive dumpers across their entire trajectory (B). The half violin plot shows the distribution of bootstrapped differences of mean gaze angular velocity of experienced and naive dumpers (C). In (B) and (C), the solid dot shows mean, while the vertical bar shows 95% confidence interval of the mean.

## DISCUSSION

In summary, our displacement study on the dumpers of the Australian red honey ant *Melophorus bagoti* revealed significant differences in navigational capability between naive and experienced dumpers. The results supported our predictions in the main. Naive dumpers as a group were not oriented towards the nest from 2 m away in any direction, whereas experienced dumpers demonstrated nestward orientation from all test locations. Naive dumpers as a group, however, were significantly nestward oriented at the test location nearest their dumping location, supporting our prediction and suggesting that they had learned something of the views around their dumping site. Furthermore, at the displacement sites, naive dumpers exhibited more meandering behaviour, characterized by increased sinuosity and reduced straightness compared to experienced dumpers. Additionally, naive dumpers walked at a slower pace compared to the experienced dumpers. These findings suggest that naïve dumpers have less navigational knowledge compared with experienced dumpers, as a result of which they scanned their surroundings more frequently and for a longer duration than did experienced dumpers. Overall, the naive dumpers lacked full knowledge of the visual cues around their nest.

The differences in travel characteristics between experienced and naive dumpers in this study parallel differences found on the dumping job and described in our recent study (Deeti et al., submitted b). On the job too, the naive dumpers walk slower and more sinuously, taking more bouts of scanning. We interpreted these characteristics as supporting learning on the job, learning the visual surround for returning home. We would interpret differences found on displacement tests in this study similarly: naive dumpers, on encountering what was likely an unfamiliar scene at most of the displacement sites, engaged in behaviours conducive to learning the visual surrounds, including stopping to look around.

Although not a main focus here, we did find in this study that a small number of naive dumpers took one or two learning walks before taking on the dumping task. These observations show that a lack of learning walks before dumping is not obligatory. Ants show some flexibility in this aspect of the job. If dumpers learn much like would-be foragers learn (Deeti and Cheng, 2021), then one learning walk would mean that they can orient homeward from 2-m distance from the nest on tests. The preference in this study and in the study on travel characteristics on the dumping job (Deeti et al., submitted b) suggests that from the nest’s perspective, dumping waste is urgent business. As before, we suggest that from the nest’s perspective, it is better not to wait the few minutes for a dumper to take learning walks, but rather, it pays to have the dumper take the waste as soon as possible even if it means having to travel slower and to learn on the job. As before (Deeti et al., submitted b), we maintain that more investigation is needed to confirm such a preliminary interpretation.

This learning-on-the-job approach of naive dumpers contrasts with that of excavators, which take out stuff with far less pathogenic potential, namely sand (Deeti et al., submitted b). The excavators always took a single learning walk before engaging in excavating, even though the distances of travel are a good deal smaller than those of dumpers, ∼15–20 cm vs. metres. As reviewed in the Introduction, in the case of would-be foragers, they presumably benefit from taking multiple pre-job learning walks, in this species and in other species as well (*M. bagoti*: Deeti et al., 2021a; *C. fortis* and *noda*: Fleischmann et al., 2016; Fleischmann et al., 2018; *M. croslandi*: Jayatilaka et al., 2018; reviews: Freas et al., 2019; Zeil and Fleischmann, 2019). These multiple walks further expand navigational knowledge. Multiple learning walks up to 1 m distance in all directions from the nest entrance allowed them to orient nestwards from 10 m distance (*Cataglyphis*: Fleischmann et al. 2018; *Melophorus*: Wystrach et al. 2012; Deeti et al. 2020), in essence increasing the catchment range for the ants.

Across our two studies on the navigational behaviour of *M. bagoti* dumpers, together with considering other studies, we have the following tentative picture of the triggering cues and behaviours elicited in dumping. In eliciting cues, chemicals that emanate from dead arthropods, which comprises the waste that *M. bagoti* dumpers carry from their nest, probably trigger dumping behaviour in these workers. The chemicals that have been shown experimentally to trigger removal behaviour in ants are oleic acid (*Pogonomyrmex badius* and *Solepnopsis saevissima*: Wilson et al., 1958; *Myrmeica vindex*: Haskins and Haskins, 1974; *Myrmica rubra*: Diez et al., 2013; *Solepnopsis invicta*: Qiu et al., 2015) and linoleic acid (*M. rubra*: Diez et al., 2015; *S. invicta*: Qiu et al., 2015; review: Sun et al., 2018), thus a monounsaturated and a polyunsaturated fatty acid, respectively. Both these fatty acids increase in a corpse with time since death (Diez et al., 2013). Other cues could add potency in triggering removal behaviour, such as infection in the corpse; compared with corpses that have died from being frozen, corpses that died from fungal infection elicit stronger behaviours (*M. rubra*: Diez et al., 2015). Fungal infection, however, results in more oleic and linoleic acid in the corpse one day after death (*S. invicta*: Qiu et al., 2015), so that this additional cue might be capitalising on the fatty-acid triggering route. Waste removal behaviour is enhanced when a colony has brood, so that other signals than those from fatty acids likely also act as modulators or cues for waste removal (*M. rubra*: Pereira et al., 2020). One gene has been identified as crucial for such a triggering effect of oleic and linoleic acids: the chemosensory protein gene *Si-CSP1* (Qiu and Cheng, 2017).

Whatever the combination of cues that drive behaviour at the input end, what is output is likely to encompass both a quantitative signal and flexibility depending on context. In our study species, *M. bagoti*, waste items of higher pathogenic potential are taken farther away from the nest to be dumped (Deeti et al., 2023). In *M. rubra*, an infected corpse, compared with a corpse that died from freezing, elicits more of a range of behaviours, including moving the corpse, cleaning the nest, and self-grooming. Vertebrate animals are theorised to possess some common representation of quantities, which could apply to spatial, temporal, numerical, and perhaps other quantities (Cheng et al., 1996; Walsh, 2003). Evidence suggests that ants encode at least numerical quantities (*Myrmecina nipponica*: Cronin, 2014). We surmise that they also encode dose-dependent quantitative measures that could be translated into various behavioural outputs such as amount of nest cleaning or the distance to transport an item for dumping. The behavioural output, however, is also context dependent. In *P. badius*, a piece of filter paper dosed with oleic acid might be taken into the nest when most of the nest is mostly engaged in foraging, or dumped outside if most of the nest is in ‘cleaning mode’ (Gordon, 1983). In our study species, naive dumpers dump waste at shorter distances than do experienced dumpers (Deeti et al., submitted b), presumably factoring in their own inexperience.

The central complex of the insect brain is crucial for coordinating behaviours in navigtaion (Heinze, 2017; Steinbeck et al., 2020). We suspect that this neural region plays a crucial role in integrating inputs and generating outputs in dumping as well. A vector of some magnitude is likely an output. In both *M. rubra* (Diez et al., 2012) and *M. bagoti* (Deeti et al., submitted b), the heading direction of each ant is idiosyncratic, making a uniform distribution across the dumping population. This suggests that a code of distance determines how far to take the waste in a favoured sector. Between the inputs that trigger expression of the *Si-CSP1* gene and the vector output, neurobiologists have much to investigate to trace the connections.

In conclusion, our displacement study on the Australian red honey ant *Melophorus bagoti* highlights significant differences between naive and experienced dumpers. Naive dumpers did not demonstrate orientation towards the nest from 2 m away, whereas experienced dumpers did. Furthermore, compared to experienced counterparts, naive dumpers walked slower and displayed more meandering behaviour with increased sinuosity and reduced straightness. The findings underscore the importance of learning and experience in shaping the navigational knowledge of desert ants.

## ACKNOWLEDGEMENTS

We acknowledge the traditional custodians of the land upon which this research was conducted, the Arrernte people. Their culture and customs have nurtured and sustained this land since the Dreamtime and continue to do so today. We pay our respects to their Elders past and present. We thank the Centre for Appropriate Technology at Alice Springs, Australia for letting us work on their property and providing some storage space, and the CSIRO Arid Zone Research at Alice Springs for administrative support. We are also thankful to Cody Freas, Vito Lionetti for helping us to take panoramic images.

## Author contributions

Experimental design: SD. Data collection: SD. Data analysis: SD. Writing: SD and KC.

## Ethics standards

Australia has no ethical regulations regarding work with insects. The study was non-invasive and no long-term aversive effects were found on the nests or on the individuals studied.

## Data availability

Supplementary videos, Excel file of data and R scripts are available at Open Science framework: https://osf.io/cwm46//files/.

